# Entangled radicals may explain lithium effects on hyperactivity

**DOI:** 10.1101/2021.03.19.436086

**Authors:** Hadi Zadeh-Haghighi, Christoph Simon

## Abstract

It is known that bipolar disorder and its lithium treatment involve the modulation of oxidative stress. Moreover, it has been observed that lithium’s effects are isotope-dependent. Based on these findings, here we propose that lithium exerts its effects by influencing the recombination dynamics of a naturally occurring radical pair involving oxygen. We develop a simple model inspired by the radical-pair mechanism in cryptochrome in the context of avian magnetoreception and xenon-induced anesthesia. Our model reproduces the observed isotopic dependence in the lithium treatment of hyperactivity in rats. It predicts a magnetic-field dependence of the effectiveness of lithium, which provides one potential experimental test of our hypothesis. Our findings show that Nature might harness quantum entanglement for the brain’s cognitive processes.

## Introduction

The human brain is a magnificent system with highly complex functionalities such as learning, memory, emotion, and subjective experience, each of which is *mood* dependent^1^. Everyday millions of patients all over the world take psychiatric pharmaceuticals to stabilize their mood, yet the underlying mechanisms behind these medications remain largely unknown^2^.

Bipolar disorder (BD) is a devastating mental illness which affects 2-4% of the world population^3^. As its name implies, it entails two distinct oscillating and opposing states, a state of energy and hyperactivity (the manic phase), and a state of low energy (depression)^4^. Lithium (Li) administration is the first-line treatment of bipolar illness^3,5–7^. Despite the common clinical use of this drug, the mechanism by which it exerts its effects remains elusive^8,9^. Here, based on experimental findings, we propose a mechanism for Li’s therapeutic effects on BD.

Studies in the 1980s showed that administering Li results in different parenting behaviors and potentially delayed offspring development in rats^10^. Although these findings weren’t quantitative, it was reported that different Li isotopes have different impacts. Recently, a new study was conducted by Ettenberg *et al*.^11^ demonstrating an isotope effect of Li on the manic phase in rats. Li has two stable isotopes, ^6^Li and ^7^Li, which carry different nuclear spin angular momentum, *I*_6_ = 1 and *I*_7_ = 3/2, respectively. In that study, sub-anesthetic doses of ketamine were administered to induce hyperactivity which was then treated with lithium. The findings of that work indicate that ^6^Li produces a longer suppression of hyperactivity in an animal model of mania compared to ^7^Li.

Moreover, there is accumulating evidence indicating that BD^12–19^ and its Li treatment^7,15,20,21^ are associated with oxidative stress, which is an imbalance between production and accumulation of radical oxygen species (ROS) in cells and tissues and the ability of a biological system to detoxify these reactive products, essential for governing life processes^22^. It therefore seems pertinent to explore the relationship between nuclear spin effects and oxidative stress in lithium mania therapy.

Any proposed mechanism for Li’s BD treatment should embrace these two facts: Li’s effect on BD exhibits isotopic dependence, and BD is connected to abnormalities in oxidative stress. In other words, the balance of naturally occurring free radicals is disrupted in BD patients, and lithium’s different isotopes appear to alter the free radical formations differently. It is therefore possible that nuclear spin properties might be the key for the differential effects of the two lithium isotopes in BD. Here, we propose that lithium affects naturally occurring radical pairs (RPs) in a way that is modulated by the hyperfine interaction (HFI).

Spins can play crucial roles in chemical reactions even though the energies involved are orders of magnitude smaller than the thermal energy, *k*_*B*_*T* ^23,24^. It has been known since the 1970s that external magnetic fields and nuclear spins can alter the rates and product yields of certain chemical reactions^25,26^. The key ingredients are RPs created simultaneously, such that the two electron spins, one on each radical, are entangled. Organic radicals are typically created in singlet (S) or triplet (T) entangled states by a reaction that conserves electron spin. Any spin in the vicinity or an external magnetic field can alter the extent and timing of the S-T interchange and hence the yields of products formed spin-selectively from the S and T states^27,28^.

Over the past decades, it has been proposed that that quantum physics could help answer unsolved questions in life science^29–31^. The above-described Radical Pair Mechanism (RPM) is one of the most well-established models in quantum biology; it could give a promising explanation for how migratory birds can navigate by means of Earth’s magnetic field^32,33^. The application of RPM has begun to gain momentum in numerous fields of research^34^. Most recently, Smith *et al*., inspired by the the RPM explanation of avian magnetoreception, have shown that the RPM could play a role in xenon-induced anesthesia, which exhibits isotopic dependence^35^.

In the context of avian magnetoreception, the RPM is thought to involve the protein cryptochrome (Cry)^36^, which contains the flavin adenine dinucleotide (FAD) cofactor. It is known that in Cry RPs can be in the form of flavosemiquinone radical (FAD^•–^) and terminal tryptophan radical (TrpH^•+^)^32,37,38^. However, considerable evidence suggests that the superoxide radical, O_2_^•–^, can be an alternative partner for the flavin radical, such that FADH^•^and O_2_^•–^ act as the donor and the acceptor, respectively^39–42^. This is motivated by the fact that the magnetically sensitive step in the reaction scheme occurs in the dark^43^, with required prior exposure to light, and that a strongly asymmetric distribution of hyperfine couplings across the two radicals results in a stronger magnetic field effect than a more even distribution^44^. Furthermore, Zhang *et al*.^45^ recently showed that a magnetic field much weaker than that of the Earth attenuates adult hippocampal neurogenesis and cognition and that this effect is mediated by ROS. Additionally, it has been shown that the biological production of ROS *in vivo* can be influenced by oscillating magnetic fields at Zeeman resonance, indicating coherent S-T mixing in the ROS formation^46^.

BD is characterized by shifts in energy, activity, and mood and is correlated with disruptions in circadian rhythms^47,48^ and abnormalities in oxidative stress^12–19^. It is known that Li influences the circadian clock in humans, and circadian rhythms are disrupted in patients with BD for which Li is a common treatment^49,50^. Yet, the exact mechanisms and pathways behind this therapy are under debate. However, it has been shown that Li acts directly on the mammalian suprachiasmatic nucleus (SCN), a circadian pacemaker in the brain^51,52^. Cry’s are necessary for SCN’s development of intercellular networks that subserve coherent rhythm expression^53^. Further, it has also been reported that Cry2 is associated with BD^54^. In addition, Cry’s essential role has been discovered in axon outgrowth in low intensity repetitive transcranial magnetic stimulation (rTMS)^55^. It thus seems that Cry might also have vital roles in the brain’s functionalities related to BD. The [FADH^•^… O_2_^•–^] RP formation in Cry is consistent with evidence that Li treatment^7,15,20,21^ and BD^12–19^ are associated with oxidative stress implying a key role for radicals in BD and Li effects. Motivated by this reasoning, in the present work, we specifically focus on the Cry pathway for Li therapy, however, this is not the only way in which radical pairs could play a role in lithium effects on BD. Alternative paths will be discussed briefly in the Discussion section.

Here, we propose that the RPM could be the underlying mechanism behind the isotope effects of Li treatment for hyperactivity. We propose that Li’s nuclear spin modulates the recombination dynamics of S-T interconversion in the naturally occurring RPs in the [FADH^•^… O_2_^•–^] complex, and due to the distinct nuclear spins, each isotope of Li influences these dynamics differently, which results in different therapeutic effects, see Fig. 1.

**Figure 1.**
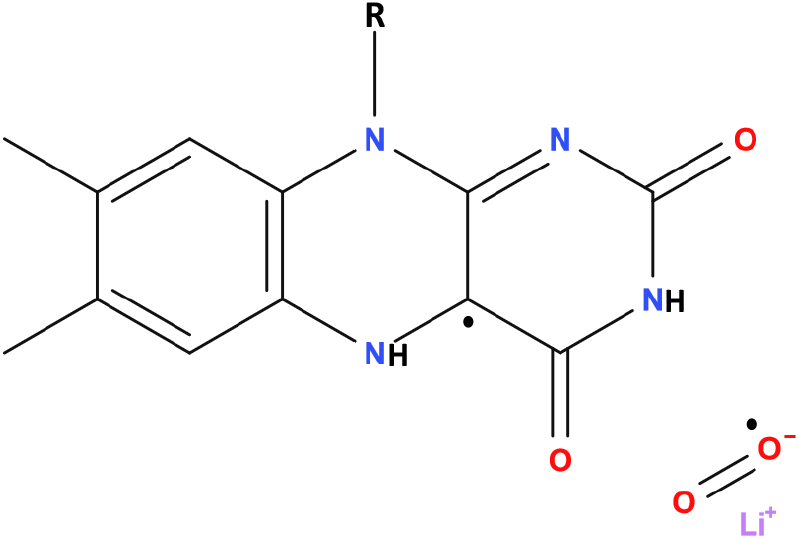
Flavinsemiquinone [FADH^•^] and lithium superoxide [Li^+^-O_2_^•−^] radical pair. The radical pair undergoes intersystem crossing between singlet and triplet entangled states.

Let us note that an alternative interpretation has also been proposed for the isotope dependence in lithium’s effect on mania. Fisher has suggested that phosphorus nuclear spins could be entangled in networks of Posner molecules, *Ca*_9_(*PO*_4_)_6_, which could form the basis of a quantum mechanism for neural processing in the brain^56^. Replacement of calcium at the center of a Posner molecule with another cation can further stabilize the Posner molecule^57^, where each isotope of the cation, due to their different nuclear spins, has a different impact. This also provides a potential interpretation for the lithium isotope dependence treatment for mania. However, this model demands more supporting evidence and has been recently challenged experimentally^58^. Moreover, our proposed explanation based on the RPM allows us to make connections with the essential roles of ROS^45^ and Cry^55^ in the brain’s neuronal activity and in the biological pathways for lithium effects on BD. In the following, we aim to probe the possibility of RPM for lithium treatment and analyze the parameter values that are required in order to explain the isotopic effect of Li’s BD treatment observed by Ettenberg *et al*.^11^.

## Results

### Lithium Effectiveness on hyperactivity and RPM

#### Quantifying effectiveness

Ettenberg *et al*.^11^ recorded the effects of lithium treatments on ketamine-induced locomotor activity of rats. They conducted the test over 60 min sessions beginning immediately after ketamine injection. After administering of ^6^Li and ^7^Li, they measured the mean traveled distance with the standard error of the mean (SEM) for every five minutes for a group of 16 rats for each isotope treatment. They observed that ^6^Li treatment exhibited significantly greater and more prolonged reductions in hyperactivity compared to ^7^Li. Here we define the cumulative traveled distance in 60 minutes *TD*_6_ and *TD*_7_, respectively, for ^6^Li and ^7^Li. The total traveled distance ratio *TD*_*r*_ is defined such that *TD*_*r*_ = *TD*_7_*/TD*_6_, see Table 1. We derived the uncertainties based on the reported mean (*±*SEM) hyperlocomotion from Ettenberg *et al*.^11^ using standard error propagation.

**Table 1.** Li isotopic nuclear spins and total traveled distance in 60 minutes for ^6^Li and ^7^Li, *TD*_6_ and *TD*_7_, respectively, taken form the work of Ettenberg *et al*.^11^. *TD*_*r*_ is the total traveled distance ratio.

| Treatment | Total Traveled Distance (cm) | Uncertainty |
| --- | --- | --- |
| <sup>6</sup>Li ($I = 1$) | $TD_6 = 20112$ | $\pm 4.62\%$ |
| <sup>7</sup>Li ($I = 3/2$) | $TD_7 = 25630$ | $\pm 3.83\%$ |
| $TD_r = TD_7/TD_6 = 1.3 \pm 6\%$ | | |

#### RPM model

The RP model developed here is used to reproduce the effectiveness of Li for treatment of hyperactivity based on alteration in the triplet yield for different isotopes. Here, we propose that Li may interact with the RP system of FADH and superoxide. The correlated spins of RP are assumed to be in [FADH^•^… O_2_^•–^], where the unpaired electron on O_2_^•–^ couples with the Li nuclear spin, see Fig. 1.

In this model, we consider a simplified form of interactions for the RPM by only including Zeeman and HF interactions^32, 59^. Given the likely randomized orientation of the relevant proteins, for the HFIs, we only consider the isotropic Fermi contact contributions. In our calculations, we assume that the unpaired electron on FADH^•^ couples only with the isoalloxazine nitrogen nucleus, which has the largest isotropic HF coupling constant (HFCC) among all the atoms in FAD^60^, following the work of Hore^61^, and that the other unpaired electron on [Li^+^-O_2_^•–^] couples with the lithium nucleus. The Hamiltonian for our RP system reads as follows:

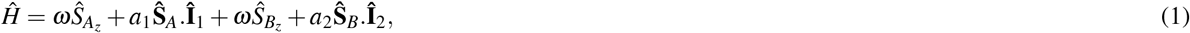

where **Ŝ**_*A*_ and **Ŝ**_*B*_ are the spin operators of radical electron A and B, respectively, **Î**_1_ is the nuclear spin operator of the isoalloxazine nitrogen of FADH^•^, **Î**_2_ is the nuclear spin operator of the Li nucleus, *a*_1_ is the HFCC between the isoalloxazine nitrogen of FADH^•^and the radical electron A (*a*_1_ = 523.3*µT* ^60^), *a*_2_ is the HFCC between the Li nucleus and the radical electron B, and *ω* is the Larmor precession frequency of the electrons due to the Zeeman effect.

#### DFT

We use density functional theory (DFT) to determine the reasonable range for the Li HFCC. Our DFT calculations show that the unpaired electron of [Li^+^-O_2_^•–^] is bound. The the highest occupied molecular orbital (HOMO) is shown in Fig. 2. The resulting Mulliken charge and spin population of the [Li^+^-O_2_^•–^] complex indicates that the unpaired electron resides primarily on the O_2_ molecule but is extended slightly onto the lithium atom, see Table 2.

**Table 2.** Mulliken charge and spin population of [Li^+^… O_2_^•−^].

| Atom | Charge Population | Spin Population |
| --- | --- | --- |
| O | -0.244409 | 0.524734 |
| O | -0.244398 | 0.524744 |
| Li | 0.488807 | -0.049479 |
| Sum | 0 | 1 |

**Figure 2.**
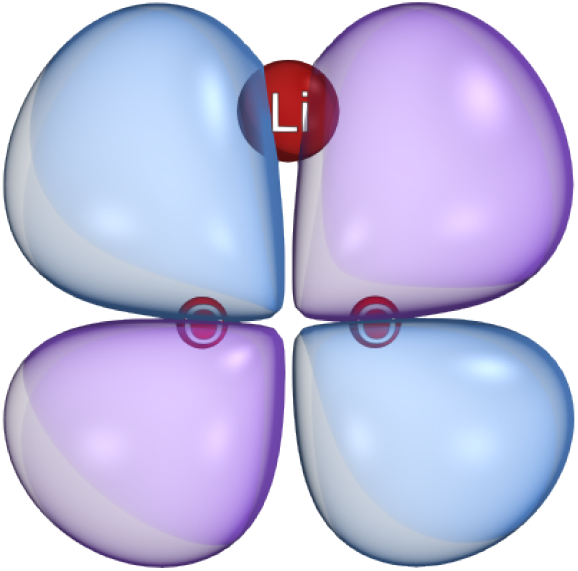
The highest occupied molecular orbital of [Li^+^-O_2_^•−^]. Imaged rendered using IboView v20150427 2 (http://www.iboview.org).

Using different DFT functionals and basis-sets, we find a range of values for the HFCC of Li nucleus at the distance of *∼*1.6Å from O_2_^•–^, which is close to the values from other studies^62^, *a*_2_ ∈ [157.7, 282]*µ*T. The HFCC values between the unpaired electron on FADH^•^ and the isoalloxazine nitrogen nuclear spin are taken from Maeda *et al*.^60^.

#### Triplet Yield Calculation

The triplet yield produced by the radical pair reaction can be obtained by tracking the spin state of the radical pair over the course of the reaction. This can be carried out by solving the Liouville-von Neumann equation, which describes the evolution of the density matrix over time. The eigenvalues and eigenvectors of the Hamiltonian can be used to determine the ultimate triplet yield (Φ_*T*_) for time periods much greater than the RP lifetime:

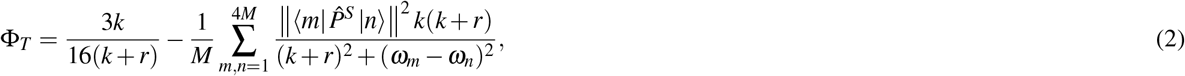

where *M* is the total number of nuclear spin configurations, 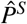 is the singlet projection operator, |*m*⟩ and |*n*⟩ are eigenstates of *Ĥ* with corresponding eigenenergies of *ω*_*m*_ and *ω*_*n*_, respectively, *k* is the RP lifetime rate, and *r* is the RP spin-coherence lifetime rate.

Here we explore the sensitivity of the triplet yield ratio to changes in the HFCC *a*_2_ between unpaired electron B and the lithium nucleus, external magnetic field strength (*B*), RP reaction rate (*k*), and RP spin-coherence relaxation rate (*r*). For the comparison between the experimental measurements and our RPM model, the absolute value of the difference between total traveled distance ratio and triplet yield, |*TD*_*r*_ −*TY*_*r*_|, is presented on the *a*_2_ and *k* plane in Fig. 3a, on the *a*_2_ and *r* plane in Fig. 3b, and on the *k* and *r* plane in Fig. 3c. The dependence of the *TY* for each isotope of lithium and their ratio on the external magnetic field is shown in Fig. 4. The experimental findings of the isotopic-dependence of Li treatment for hyperactivity are reproducible for *B* ∈ [0, 200]*µ*T, which includes the geomagnetic field at different geographic locations (25 to 65*µ*T)^63^. We aim to find regions in parameter space by which the triplet yield calculated from our RPM model fits with the experimental findings on the isotope dependence of the effectiveness of lithium for hyperactivity treatment. We indicate the regions within which the difference between our model and the experimental data is smaller than the uncertainty of the experimental results,*i*.*e*., |*TD*_*r*_ − *TY*_*r*_| ≤0.06, as shown in Fig. 3. For a fixed external magnetic field *B* = 50*µT* and *a*_1_ = 523.3*µ*T, our model can predict the experimental results with quite broad ranges for the HFCC between the unpaired electron and Li nuclear spin, *a*_2_, the RP reaction rate, *k*, the RP spin relaxation rate, *r*, namely *a*_2_ ∈ [210, 500]*µ*T, *k* ∈ [1. × 10^8^, 1× 10^9^] s^−1^, and *r* ∈ [1 × 10^6^, 6 × 10^7^] s^−1^, see Fig. 3. The predicted values for the lithium HFCC overlaps with the range that we find for Li^+^-O_2_^•–^ from our DFT calculations.

**Figure 3.**
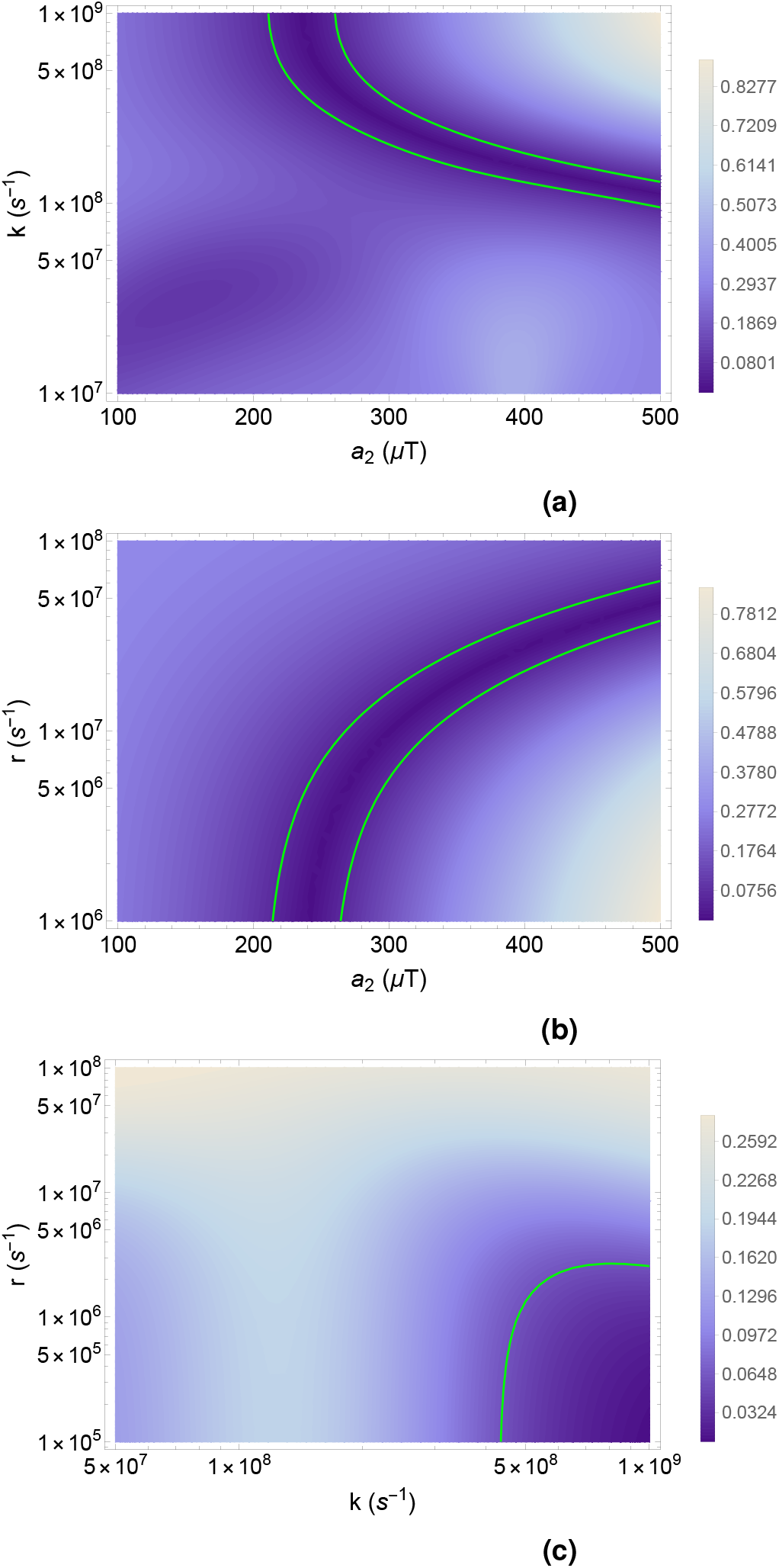
The dependence of the agreement between the total traveled distance ratio, *TD*_*r*_, and the triplet yield ratio, *TY*_*r*_ of ^7^Li over ^6^Li on the relationship between: a) the radical pair reaction rate, *k*, and the lithium hyperfine coupling constant, *a*_2_, for *r* = 1.0×10^6^ s^−1^ b) the radical pair spin-coherence relaxation rate, *r*, and the lithium hyperfine coupling constant, *a*_2_, for *k* = 7.0×10^8^ s^−1^ and c) the RP reaction rate, *k* and the radical pair spin-coherence relaxation rate, *r*, for *a*_2_ = 224.4*µ*T. In all three cases *a*_1_ = 523.3*µ*T and *B* = 50*µ*T. The absolute value of the difference between the prediction of radical pair mechanism, *TY*_*r*_, and the experimental data, *TD*_*r*_, is illustrated where the green line indicates the ranges smaller than the experimental uncertainty, |*TD*_*r*_ −*TY*_*r*_| ≤ 0.06.

**Figure 4.**
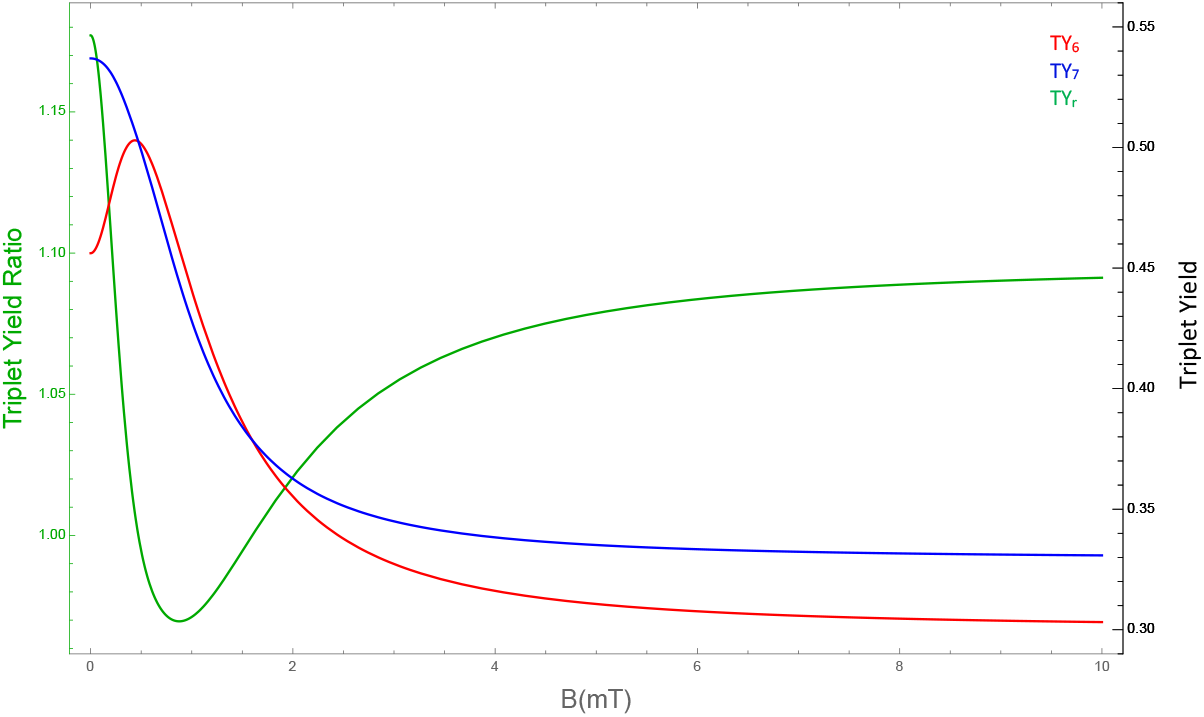
The dependence of the triplet yield of ^6^Li, *TY*_6_, (red) and ^7^Li, *TY*_7_, (blue) and the triplet yield ratio, *TY*_*r*_, (green) on external magnetic field *B* for *a*_1_ = 523.3*µ*T, *a*_2_ = 224.4*µ*T, *r* = 1.0×10^6^ s^−1^, and *k* = 4.0×10^7^ s^−1^.

## Discussion

Our principal goal in this project was to probe whether the RPM can be the underlying mechanism behind the isotopic dependence of lithium’s effectiveness for the hyperactivity treatment observed by Ettenberg *et al*.^11^. Our results support such a mechanism. A simple radical model with a set of reasonable parameters can reproduce the experimental findings.

We proposed that the [FADH^•^… O_2_^•–^] complex is the naturally occurring RP in the circadian center of the brain and lithium interacts with the radical electron on superoxide. This is motivated by the observations^51^ that one of the possible pathways for Li’s effects is Li’s direct influence on the suprachiasmatic nucleus–a region in the brain containing cryptochrome protein–, that bipolar disorder is associated with imbalances in the ROS level^14^, and that the lithium treatment modulates the oxidative stress level^7,15,20,21^. By varying the RP spin-coherence relaxation rate, RP reaction rate, and hyperfine coupling parameters our model reproduced the experimental findings of Ettenberg *et al*.^11^. The predicted range for the lithium hyperfine coupling constant in Li^+^-O_2_^•–^ overlaps with our DFT calculations.

Let us note that the RPM model could be adapted for other pathways by which lithium could affect BD. For example, it has also been proposed that Li could exert its effects on BD via increasing glutamate re-uptake at the N-methyl-D-aspartate (NMDA) receptor^64^. Inspired by the RPM’s potential role in xenon-induced anesthesia^35^, where there the NMDA receptor has been proposed as the target site of xenon, a similar scenario could be envisioned here. For example, oxygen might oxidize Trp in the NMDA receptor and result in the formation of [TrpH^•–^…O_2_^•–^] RPs, and lithium might modulate the S-T interconversion of the RPs by coupling to the unpaired electron of O_2_^•–^.

There remains a question on the feasibility for the O_2_^•–^ radical to be involved in the RPM in this scenario due to its expected fast spin relaxation rate *r*. In the context of magnetoreception, the superoxide-based RP model has been discussed by Hogben *et al*.^44^ and Player and Hore^65^. The authors argue that due to fast molecular rotation free O_2_^•–^ has a spin relaxation lifetime on the orders of 1 ns. The relaxation rate requirement calculated by our model yields *r* about two orders of magnitude slower than this value, see Fig. 3. However, the same authors have also noted that this fast spin relaxation of free superoxide can be can be lowered if the molecular symmetry is reduced and the angular momentum quenched by its biological environment. Moreover, Kattnig^66,67^ proposed that such fast spin relaxation of O_2_^•–^ could be, in effect, reduced by the involvement of scavenger species around O_2_^•–^. For example, it has been suggested that, in the RPM, Trp^•^ could act as a scavenger molecule for O_2_^•–42^. Alternatively, other RP constituents could be considered instead of O_2_^•–^ to explain isotopic effects within the framework of the RPM.

The predicted dependence of the triple yield on changes of external magnetic field in Fig.4 indicates that the effectiveness of the Li treatment could be enhanced by applying external magnetic fields. It would be of interest to conduct such experiments *in vivo* to explore the impact of the external magnetic field on the effectiveness of the different isotopes of lithium for hyperactivity treatment. It would also be of interest, to explore isotopic nuclear-spin effects of key elements of the biological environment (particularly oxygen, but also e.g. nitrogen, carbon, and hydrogen) in experiments on the lithium effectiveness on hyperactivity.

In summary, our results suggest that quantum entanglement might lie behind the mechanism of lithium treatment for BD, similarly to magnetoreception in animals^33^ and (as recently proposed) xenon-induced anesthesia^35^.

This also raises the question whether the RPM could play a role in other mental disorders, and could lead to new approaches to treatment and improving efficiency of medications^68^, specifically for illnesses that have been shown to be associated with oxidative stress, such as Alzheimer’s^69,70^, Schizophrenia^71–73^, and Parkinson’s^74^. Further, it is known that light exposure affects mood and emotions^75–80^, and light is required for the formation of RPs in Crys^39,65^. It therefore seems that RPM could also elucidate the effects of light exposure on mood. Similarly, RPM may provide explanations for the anti-depressant effect of vitamin D^81–83^ and its effects on the modulation of ROS production^84, 85^. Given that the RPM is typically associated with isotope and magnetic field effects, it would be of great interest to search for such effects for other neurological medications.

Memory, learning, and subjective experience are affected by moods and emotions, and it has been proposed that cognition and subjective experience may be related to large-scale entanglement^56, 86–89^. In this highly speculative context, entangled RPs could play crucial roles as sources of entanglement, and the present work could be viewed as another piece of evidence consistent with this idea in addition to Ref.^35^. In particular, superoxide radicals can give rise to singlet oxygen, which can emit photons^90^. These photons could serve as quantum messengers to establish long-distance connections^91^ that might be essential for consciousness^92^.

## Methods

### DFT Analysis

The ORCA package^93^ was used for our Li^+^-O_2_^•–^ DFT calculations, and the molecular structure was optimized using the dispersion-corrected PBE0 functional and def2-TZVP basis set.

The orbitals obtained from the optimization calculations were used to calculate orbital energies as well as the hyperfine coupling constant *a*_2_. Using various DFT functionals, we obtained *a*_2_ ∈ [157.7, 282]*µ*T, see Table 3. The calculation with the double hybrid functional^94^ RI-B2GP-PLYP with def2-QZVPP basis-set resulted *a*_2_ = 224.4*µ*T, which is in the range that our RPM model predicts. We used this value in our computations. Relativistic effects were treated by a scalar relativistic Hamiltonian using the zeroth-order regular approximation (ZORA)^95^. Solvent effects were considered by using the conductor-like polarizable continuum model (CPCM)^96^, with a dielectric constant of 2.

**Table 3.** HFCC *a*_2_(*µ*T) using different DFT fuctionals.

| Functional | $a_2(\mu)\text{T}$ |
| --- | --- |
| BHLYP | 282 |
| RI-B2GP-PLYP | 224.4 |
| B3LYP | 210.8 |
| B3LYP/G | 209.8 |
| PBE0 | 167.2 |
| B3PW91 | 157.7 |
| PW6B95 | 208.4 |

## Data Availability

The generated datasets and computational analysis are available from the corresponding author on reasonable request.

## Acknowledgements

The authors would like to thank Dennis Salahub, Wilten Nicola, Rishabh, Mansoor Askari, and Jordan Smith for their input, comments, and insights.The authors would like to acknowledge Compute Canada for its computing resources. This work was supported by the Natural Sciences and Engineering Research Council of Canada.

## Author contributions statement

H.ZH. performed calculations and modelling with feedback from C.S.; H.ZH. and C.S. wrote the paper; C.S. conceived and supervised the project.

## Competing Interests

The authors declare no competing interests.

## Notes

### Competing Interest Statement

The authors have declared no competing interest.

## References

1. Martínez-Arán, A. et al. Cognitive function across manic or hypomanic, depressed, and euthymic states in bipolar disorder. Am. J. Psychiatry 161, 262–270 (2004).

2. Grande, I., Berk, M., Birmaher, B. & Vieta, E. Bipolar disorder. The Lancet 387, 1561–1572 (2016).

3. Geddes, J. R. & Miklowitz, D. J. Treatment of bipolar disorder. The lancet 381, 1672–1682 (2013).

4. McIntyre, R. S. et al. Bipolar disorders. The Lancet 396, 1841–1856 (2020).

5. Malhi, G. S., Gessler, D. & Outhred, T. The use of lithium for the treatment of bipolar disorder: recommendations from clinical practice guidelines. J. affective disorders 217, 266–280 (2017).

6. Machado-Vieira, R., Manji, H. K. & Zarate Jr, C. A. The role of lithium in the treatment of bipolar disorder: convergent evidence for neurotrophic effects as a unifying hypothesis. Bipolar disorders 11, 92–109 (2009).

7. Khairova, R. et al. Effects of lithium on oxidative stress parameters in healthy subjects. Mol. Medicine Reports 5, 680–682 (2012).

8. Roux, M. & Dosseto, A. From direct to indirect lithium targets: a comprehensive review of omics data. Metallomics 9, 1326–1351 (2017).

9. Harwood, A. Lithium and bipolar mood disorder: the inositol-depletion hypothesis revisited. Mol. psychiatry 10, 117–126 (2005).

10. Sechzer, J. A., Lieberman, K. W., Alexander, G. J., Weidman, D. & Stokes, P. E. Aberrant parenting and delayed offspring development in rats exposed to lithium. Biol. psychiatry 21, 1258–1266 (1986).

11. Ettenberg, A. et al. Differential effects of lithium isotopes in a ketamine-induced hyperactivity model of mania. Pharmacol. Biochem. Behav. 190, 172875 (2020).

12. Andreazza, A. C. et al. Oxidative stress markers in bipolar disorder: a meta-analysis. J. affective disorders 111, 135–144 (2008).

13. Yumru, M. et al. Oxidative imbalance in bipolar disorder subtypes: a comparative study. Prog. Neuro-Psychopharmacology Biol. Psychiatry 33, 1070–1074 (2009).

14. Salim, S. Oxidative stress and psychological disorders. Curr. neuropharmacology 12, 140–147 (2014).

15. Machado-Vieira, R. et al. Oxidative stress parameters in unmedicated and treated bipolar subjects during initial manic episode: a possible role for lithium antioxidant effects. Neurosci. letters 421, 33–36 (2007).

16. Ng, F., Berk, M., Dean, O. & Bush, A. I. Oxidative stress in psychiatric disorders: evidence base and therapeutic implications. Int. J. Neuropsychopharmacol. 11, 851–876 (2008).

17. Lee, S.-Y. et al. Oxidative/nitrosative stress and antidepressants: targets for novel antidepressants. Prog. Neuro-Psychopharmacology Biol. Psychiatry 46, 224–235 (2013).

18. Brown, N. C., Andreazza, A. C. & Young, L. T. An updated meta-analysis of oxidative stress markers in bipolar disorder. Psychiatry Res. 218, 61–68 (2014).

19. Berk, M. et al. Pathways underlying neuroprogression in bipolar disorder: focus on inflammation, oxidative stress and neurotrophic factors. Neurosci. & biobehavioral reviews 35, 804–817 (2011).

20. Frey, B. N. et al. Effects of lithium and valproate on amphetamine-induced oxidative stress generation in an animal model of mania. J. Psychiatry Neurosci. 31, 326 (2006).

21. de Sousa, R. T. et al. Oxidative stress in early stage bipolar disorder and the association with response to lithium. J. psychiatric research 50, 36–41 (2014).

22. Sies, H., Berndt, C. & Jones, D. P. Oxidative stress. Annu. review biochemistry 86, 715–748 (2017).

23. Steiner, U. E. & Ulrich, T. Magnetic field effects in chemical kinetics and related phenomena. Chem. Rev. 89, 51–147 (1989).

24. Hayashi, H. Introduction to dynamic spin chemistry: magnetic field effects on chemical and biochemical reactions, vol. 8 (World Scientific Publishing Company, 2004).

25. Schulten, K., Staerk, H., Weller, A., Werner, H.-J. & Nickel, B. Magnetic field dependence of the geminate recombination of radical ion pairs in polar solvents. Z. Phys. Chem 101, 371–390 (1976).

26. Brocklehurst, B. et al. The effect of a magnetic field on the singlet/triplet ratio in geminate ion recombination. Chem. Phys. Lett. 28, 361–363 (1974).

27. Jones, A. R. Magnetic field effects in proteins. Mol. Phys. 114, 1691–1702 (2016).

28. Timmel, C. R., Till, U., Brocklehurst, B., Mclauchlan, K.A. & Hore, P.J. Effects of weak magnetic fields on free radical recombination reactions. Mol. Phys. 95, 71–89 (1998).

29. Ball, P. Physics of life: The dawn of quantum biology. Nat. News 474, 272–274 (2011).

30. McFadden, J. & Al-Khalili, J. Life on the edge: the coming of age of quantum biology (Crown Publishing Group (NY), 2016).

31. Kim, Y. et al. Quantum biology: An update and perspective. Quantum Reports 3, 80–126 (2021).

32. Hore, P. J. & Mouritsen, H. The radical-pair mechanism of magnetoreception. Annu. review biophysics 45, 299–344 (2016).

33. Mouritsen, H. Long-distance navigation and magnetoreception in migratory animals. Nature 558, 50–59 (2018).

34. Hore, P. J., Ivanov, K. L. & Wasielewski, M. R. Spin chemistry. The J. Chem. Phys. 152, 120401 (2020).

35. Smith, J., Zadeh-Haghighi, H., Salahub, D. & Simon, C. Radical pairs may play a role in xenon-induced general anesthesia. Sci. Reports 11, 6287 (2021).

36. Ritz, T., Adem, S. & Schulten, K. A model for photoreceptor-based magnetoreception in birds. Biophys. journal 78, 707–718 (2000).

37. Giovani, B., Byrdin, M., Ahmad, M. & Brettel, K. Light-induced electron transfer in a cryptochrome blue-light photoreceptor. Nat. Struct. & Mol. Biol. 10, 489–490 (2003).

38. Hong, G., Pachter, R., Essen, L.-O. & Ritz, T. Electron transfer and spin dynamics of the radical-pair in the cryptochrome from chlamydomonas reinhardtii by computational analysis. The J. chemical physics 152, 065101 (2020).

39. Müller, P. & Ahmad, M. Light-activated cryptochrome reacts with molecular oxygen to form a flavin–superoxide radical pair consistent with magnetoreception. J. Biol. Chem. 286, 21033–21040 (2011).

40. Romero, E., Gómez Castellanos, J.R., Gadda, G., Fraaije, M. W. & Mattevi, A. Same substrate, many reactions: Oxygen activation in flavoenzymes. Chem. reviews 118, 1742–1769 (2018).

41. Chaiyen, P., Fraaije, M. W. & Mattevi, A. The enigmatic reaction of flavins with oxygen. Trends biochemical sciences 37, 373–380 (2012).

42. Mondal, P. & Huix-Rotllant, M. Theoretical insights into the formation and stability of radical oxygen species in cryptochromes. Phys. Chem. Chem. Phys. 21, 8874–8882 (2019).

43. Wiltschko, R., Ahmad, M., Nießner, C., Gehring, D. & Wiltschko, W. Light-dependent magnetoreception in birds: the crucial step occurs in the dark. J. The Royal Soc. Interface 13, 20151010 (2016).

44. Hogben, H. J., Efimova, O., Wagner-Rundell, N., Timmel, C. R. & Hore, P. Possible involvement of superoxide and dioxygen with cryptochrome in avian magnetoreception: origin of zeeman resonances observed by in vivo epr spectroscopy. Chem. Phys. Lett. 480, 118–122 (2009).

45. Zhang, B. et al. Long-term exposure to a hypomagnetic field attenuates adult hippocampal neurogenesis and cognition. Nat. communications 12, 1–17 (2021).

46. Usselman, R. J. et al. The quantum biology of reactive oxygen species partitioning impacts cellular bioenergetics. Sci. reports 6, 1–6 (2016).

47. Abreu, T. & Bragança, M. The bipolarity of light and dark: a review on bipolar disorder and circadian cycles. J. affective disorders 185, 219–229 (2015).

48. Takahashi, J. S., Hong, H.-K., Ko, C. H. & McDearmon, E. L. The genetics of mammalian circadian order and disorder: implications for physiology and disease. Nat. reviews genetics 9, 764–775 (2008).

49. Yin, L., Wang, J., Klein, P. S. & Lazar, M. A. Nuclear receptor rev-erbα is a critical lithium-sensitive component of the circadian clock. Science 311, 1002–1005 (2006).

50. Li, J., Lu, W.-Q., Beesley, S., Loudon, A. S. & Meng, Q.-J. Lithium impacts on the amplitude and period of the molecular circadian clockwork. PloS one 7, e33292 (2012).

51. Abe, M., Herzog, E. D. & Block, G. D. Lithium lengthens the circadian period of individual suprachiasmatic nucleus neurons. Neuroreport 11, 3261–3264 (2000).

52. Osland, T. M. et al. Lithium differentially affects clock gene expression in serum-shocked nih-3t3 cells. J. psychopharmacology 25, 924–933 (2011).

53. Welsh, D. K., Takahashi, J. S. & Kay, S. A. Suprachiasmatic nucleus: cell autonomy and network properties. Annu. review physiology 72, 551–577 (2010).

54. Lavebratt, C. et al. Cry2 is associated with depression. Plos one 5, e9407 (2010).

55. Dufor, T. et al. Neural circuit repair by low-intensity magnetic stimulation requires cellular magnetoreceptors and specific stimulation patterns. Sci. advances 5, eaav9847 (2019).

56. Fisher, M. P. Quantum cognition: The possibility of processing with nuclear spins in the brain. Annals Phys. 362, 593–602 (2015).

57. Swift, M. W., Van de Walle, C. G. & Fisher, M. P. Posner molecules: from atomic structure to nuclear spins. Phys. Chem. Chem. Phys. 20, 12373–12380 (2018).

58. Chen, R., Li, N., Qian, H., Zhao, R.-H. & Zhang, S.-H. Experimental evidence refuting the assumption of phosphorus-31 nuclear-spin entanglement-mediated consciousness. J. Integr. Neurosci. 19, 595–600 (2020).

59. Efimova, O. & Hore, P. Role of exchange and dipolar interactions in the radical pair model of the avian magnetic compass. Biophys. J. 94, 1565–1574 (2008).

60. Maeda, K. et al. Magnetically sensitive light-induced reactions in cryptochrome are consistent with its proposed role as a magnetoreceptor. Proc. Natl. Acad. Sci. 109, 4774–4779 (2012).

61. Hore, P. J. Upper bound on the biological effects of 50/60 hz magnetic fields mediated by radical pairs. Elife 8, e44179 (2019).

62. Lau, K. C., Curtiss, L. A. & Greeley, J. Density functional investigation of the thermodynamic stability of lithium oxide bulk crystalline structures as a function of oxygen pressure. The J. Phys. Chem. C 115, 23625–23633 (2011).

63. Finlay, C. C. et al. International geomagnetic reference field: the eleventh generation. Geophys. J. Int. 183, 1216–1230 (2010).

64. Ghasemi, M. & Dehpour, A. R. The nmda receptor/nitric oxide pathway: a target for the therapeutic and toxic effects of lithium. Trends pharmacological sciences 32, 420–434 (2011).

65. Player, T. C. & Hore, P. Viability of superoxide-containing radical pairs as magnetoreceptors. The J. chemical physics 151, 225101 (2019).

66. Kattnig, D. R. Radical-pair-based magnetoreception amplified by radical scavenging: resilience to spin relaxation. The J. Phys. Chem. B 121, 10215–10227 (2017).

67. Kattnig, D. R. & Hore, P. The sensitivity of a radical pair compass magnetoreceptor can be significantly amplified by radical scavengers. Sci. reports 7, 1–12 (2017).

68. Raz, A. Perspectives on the efficacy of antidepressants for child and adolescent depression. PLoS Med 3, e9 (2005).

69. Forlenza, O. V., de Paula, V. J., Machado-Vieira, R., Diniz, B. S. & Gattaz, W. F. Does lithium prevent alzheimer’s disease? Drugs & aging 29, 335–342 (2012).

70. Forlenza, O. V.,De-Paula, V.d.J.R. & Diniz, B. Neuroprotective effects of lithium: implications for the treatment of alzheimer’s disease and related neurodegenerative disorders. ACS chemical neuroscience 5, 443–450 (2014).

71. Mahadik, S. P. & Mukherjee, S. Free radical pathology and antioxidant defense in schizophrenia: a review. Schizophr. research 19, 1–17 (1996).

72. Clay, H. B., Sillivan, S. & Konradi, C. Mitochondrial dysfunction and pathology in bipolar disorder and schizophrenia. Int. J. Dev. Neurosci. 29, 311–324 (2011).

73. Gao, W.-J. & Snyder, M. A. Nmda hypofunction as a convergence point for progression and symptoms of schizophrenia. Front. cellular neuroscience 7, 31 (2013).

74. Jenner, P. Oxidative stress in parkinson’s disease. Annals Neurol. Off. J. Am. Neurol. Assoc. Child Neurol. Soc. 53, S26–S38 (2003).

75. Kent, S. T. et al. Effect of sunlight exposure on cognitive function among depressed and non-depressed participants: a regards cross-sectional study. Environ. Heal. 8, 1–14 (2009).

76. LeGates, T. A., Fernandez, D. C. & Hattar, S. Light as a central modulator of circadian rhythms, sleep and affect. Nat. Rev. Neurosci. 15, 443–454 (2014).

77. Srinivasan, V. et al. Melatonin in mood disorders. The World J. Biol. Psychiatry 7, 138–151 (2006).

78. McColl, S. L. & Veitch, J. A. Full-spectrum fluorescent lighting: a review of its effects on physiology and health. Psychol. medicine 31, 949–964 (2001).

79. Pail, G. et al. Bright-light therapy in the treatment of mood disorders. Neuropsychobiology 64, 152–162 (2011).

80. Winkler, D., Pjrek, E., Iwaki, R. & Kasper, S. Treatment of seasonal affective disorder. Expert. review neurotherapeutics 6, 1039–1048 (2006).

81. Penckofer, S., Kouba, J., Byrn, M. & Estwing Ferrans, C. Vitamin d and depression: where is all the sunshine? Issues mental health nursing 31, 385–393 (2010).

82. Spedding, S. Vitamin d and depression: a systematic review and meta-analysis comparing studies with and without biological flaws. Nutrients 6, 1501–1518 (2014).

83. Parker, G. B., Brotchie, H. & Graham, R. K. Vitamin d and depression. J. affective disorders 208, 56–61 (2017).

84. Uberti, F. et al. Protective effects of vitamin d 3 on fimbrial cells exposed to catalytic iron damage. J. ovarian research 9, 1–10 (2016).

85. Uberti, F. et al. Biological effects of combined resveratrol and vitamin d3 on ovarian tissue. J. ovarian research 10, 1–14 (2017).

86. Hameroff, S. & Penrose, R. Consciousness in the universe: A review of the ‘orch or’theory. Phys. life reviews 11, 39–78 (2014).

87. Hameroff, S. R., Craddock, T. J. & Tuszynski, J. A. Quantum effects in the understanding of consciousness. J. integrative neuroscience 13, 229–252 (2014).

88. Simon, C. Can quantum physics help solve the hard problem of consciousness? J. Conscious. Stud. 26, 204–218 (2019).

89. Adams, B. & Petruccione, F. Quantum effects in the brain: A review. AVS Quantum Sci. 2, 022901 (2020).

90. Cifra, M. & Pospíšil, P. Ultra-weak photon emission from biological samples: definition, mechanisms, properties, detection and applications. J. Photochem. Photobiol. B: Biol. 139, 2–10 (2014).

91. Kumar, S., Boone, K., Tuszyński, J., Barclay, P. & Simon, C. Possible existence of optical communication channels in the brain. Sci. reports 6, 36508 (2016).

92. Berkovitch, L. et al. Disruption of conscious access in psychosis is associated with altered structural brain connectivity. J. Neurosci. 41, 513–523 (2021).

93. Neese, F. The orca program system. Wiley Interdiscip. Rev. Comput. Mol. Sci. 2, 73–78 (2012).

94. Goerigk, L. & Grimme, S. A thorough benchmark of density functional methods for general main group thermochemistry, kinetics, and noncovalent interactions. Phys. Chem. Chem. Phys. 13, 6670–6688 (2011).

95. Van Lenthe, E. v., Snijders, J. & Baerends, E. The zero-order regular approximation for relativistic effects: The effect of spin–orbit coupling in closed shell molecules. The J. chemical physics 105, 6505–6516 (1996).

96. Marenich, A. V., Cramer, C. J. & Truhlar, D. G. Universal solvation model based on solute electron density and on a continuum model of the solvent defined by the bulk dielectric constant and atomic surface tensions. The J. Phys. Chem. B 113, 6378–6396 (2009).

